# Optimisation-based modelling for drug discovery in malaria

**DOI:** 10.1101/2022.02.12.479469

**Authors:** Yutong Li, Jonathan Cardoso-Silva, Lazaros G. Papageorgiou, Sophia Tsoka

## Abstract

The discovery of new antimalarial medicines with novel mechanisms of action is important, given the ability of parasites to develop resistance to current treatments. Through the Open Source Malaria project that aims to discover new medications for malaria, several series of compounds have been obtained and tested. Analysis of the effective fragments in these compounds is important in order to derive means of optimal drug design and improve the relevant pharmaceutical application. We have previously reported a novel optimisation-based method for quantitative structure-activity relationship modelling, modSAR, that provides explainable modelling of ligand activity through a mathematical programming formulation. Briefly, modSAR clusters small molecules according to chemical similarity, determines the optimal split of each cluster into appropriate regions, and derives piecewise linear regression equations to predict the inhibitory effect of small molecules. Here, we report application of modSAR in the analysis of OSM anti-malarial compounds and illustrate how rules generated by the model can provide interpretable results for the contribution of individual ECFP fingerprints in predicting ligand activity, and contribute to the search for effective drug treatments.

## Introduction

Malaria is an important public health problem worldwide. According to World Health Organisation,^1^ an estimated 229 million malaria cases existed in 87 malaria-endemic countries across the world, resulting in an estimated 409,000 deaths in 2019. This mosquito-borne infectious disease, caused by the *Plasmodium* parasite, affects humans and animals. Despite several approved antimalarial drugs, parasites become increasingly resistant to treatment,^2,3^ necessitating the search for new combinations of existing treatments^4–6^ or novel drugs that counteract parasite resistance.^7–10^

The development of anti-malarial drugs, however, is a complex and expensive process comprising multiple stages of compound screening and validation. Machine learning strategies can reduce the cost and time requirements associated to drug discovery by excluding unsuitable compounds and directing the search towards the most promising drug candidates.^11,12^ A typical computational strategy comprises appropriate algorithmic development and virtual screening leading to several promising candidates, which can then be analysed and further optimised for drug development.

Quantitative Structure-Activity Relationship (QSAR)^13^ modelling is also a popular technique in antimalarial drug research.^14^ In general, QSAR methods are mathematical models that aim to predict the biological activity of chemical compounds on the basis of their structural properties, and therefore can relate different functional groups of a compound to the relevant activity.^15^ In antimalarial drug research, a wide range of QSAR applications exist, for example screening a natural product library and prioritising potent antimalarial drugs, ^16^ and a QSAR model together with docking studies to find potential inhibitors for malarial resistance.^17^

Among important contributions to the development of anti-malarial drugs, the Open Source Malaria (OSM) project was established^18^ to evaluate the properties of compounds from high-throughput screens by pharmaceutical companies. Recently, a competition was launched to develop and evaluate strategies for accurate prediction of anti-PfATP4 activity among Series 4 compounds from the OSM master chemical list database, thereby reducing project costs associated with the unnecessary synthesis of inactive compounds.^19^

In this study, we build upon previous work on development of an interpretable model for QSAR modelling based on mathematical optimisation, modSAR,^20^ and model the inhibition activity (*pIC*_50_) of compounds from the OSM dataset against *Plasmodium falciparum*. We have previously applied modSAR to an earlier version of the OSM data using pre-defined molecular descriptors,^19^ and here we develop this work further in order to offer a better understanding of these antimalarial candidates as well as paving the way for future SAR explorations, lead optimisation and new *de novo* drug design efforts for malaria.

## Methodology

A schematic overview our methodology is shown in Figure 1 and comprises data pre-processing, QSAR modelling via modSAR and analysis of the rules generated by the model. ModSAR involves first detecting clusters of chemical compounds, and then applying mathematical optimisation to determine the optimal split of each cluster into appropriate regions and yield piecewise linear regression equations to link molecular descriptors to the biological activity of samples in that region. Therefore, modSAR identifies the relationship between compound features and the relevant bioactivity, in a manner that is mathematically descriptive, and with similar accuracy as those of popular machine learning methods.^20,21^ Below, data and methodology are described in more detail.

**Figure 1:**
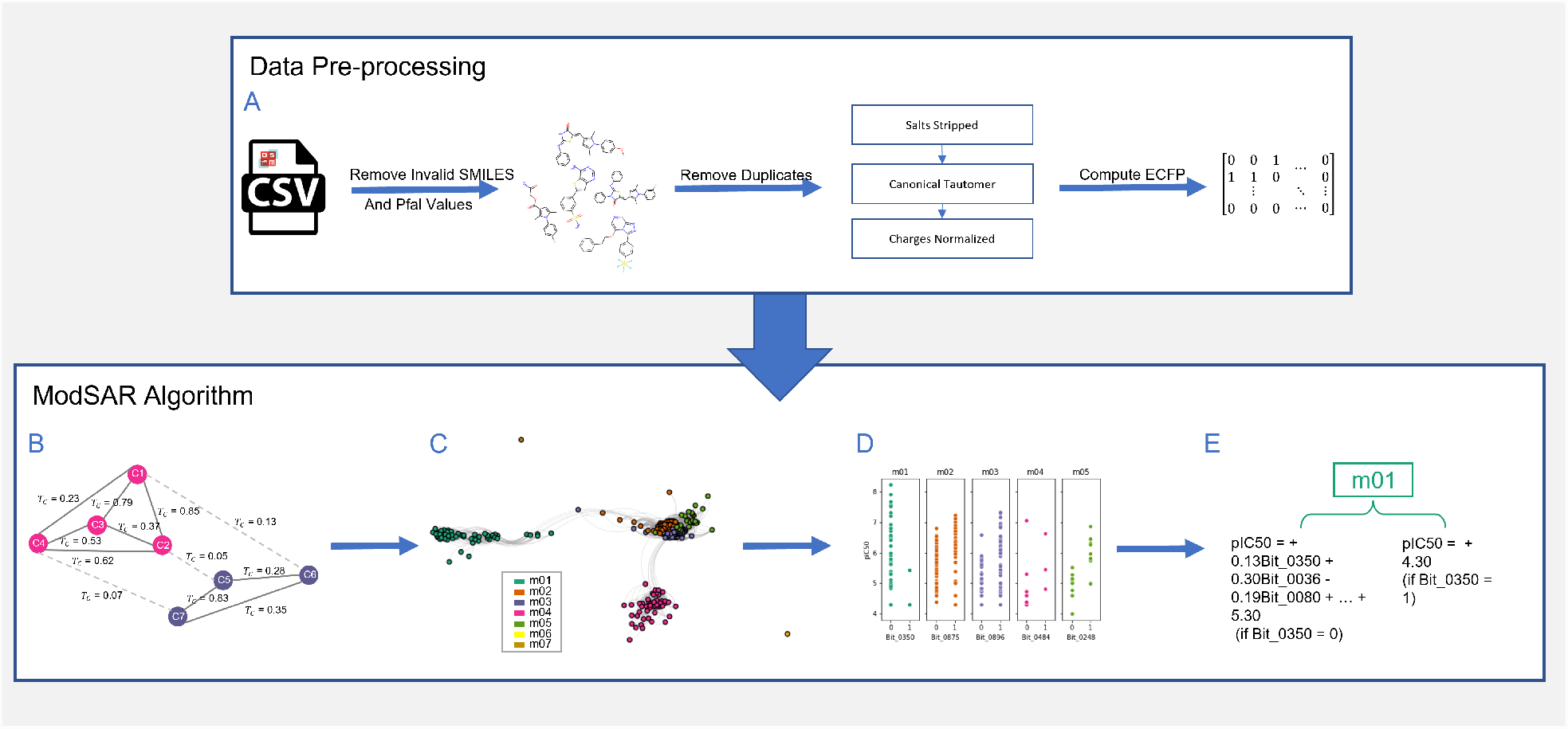
The pipeline of data processing and analysis via modSAR. (A) The OSM dataset was pre-processed following the steps specified in https://github.com/OpenSourceMalaria/Series4_PredictiveModel/issues/1, and in the end the ECFP of each OSM compound was obtained. (B) The *T*_*c*_ similarity between compounds was computed using ECFP, and a representative graph was obtained by linking compounds with respect to their similarity. (C) A threshold of similarity was applied to the graph, and different modules were identified. (D) For each module, a bit of ECFP was selected as breakpoint. Each module was then subdivided into two regions according to different values of the breakpoint feature. (E) A regression model was identified for each region.

### Data

Data used in this study derive from the Open Source Malaria (OSM) project, a collaborative consortium aiming to facilitate design of new drugs for malaria guided by open source principles,^18^ and describe the inhibitory activity (*IC*_50_) of compounds targeting *Plasmodium falciparum*. The data was downloaded from https://docs.google.com/spreadsheets/d/1Rvy6OiM291d1GN_cyT6eSw_C3lSuJ1jaR7AJa8hgGsc/edit#gid=510297618. Molecules have been categorised in four series according to chemotype: an arylpyrrole series (Series 1), the triazolourea singleton (Series 2), aminothienopyrimidine Series (Series 3), and triazolopyrazine series (Series 4). The compounds that characterise each series are shown in Figure 2.

**Figure 2:**
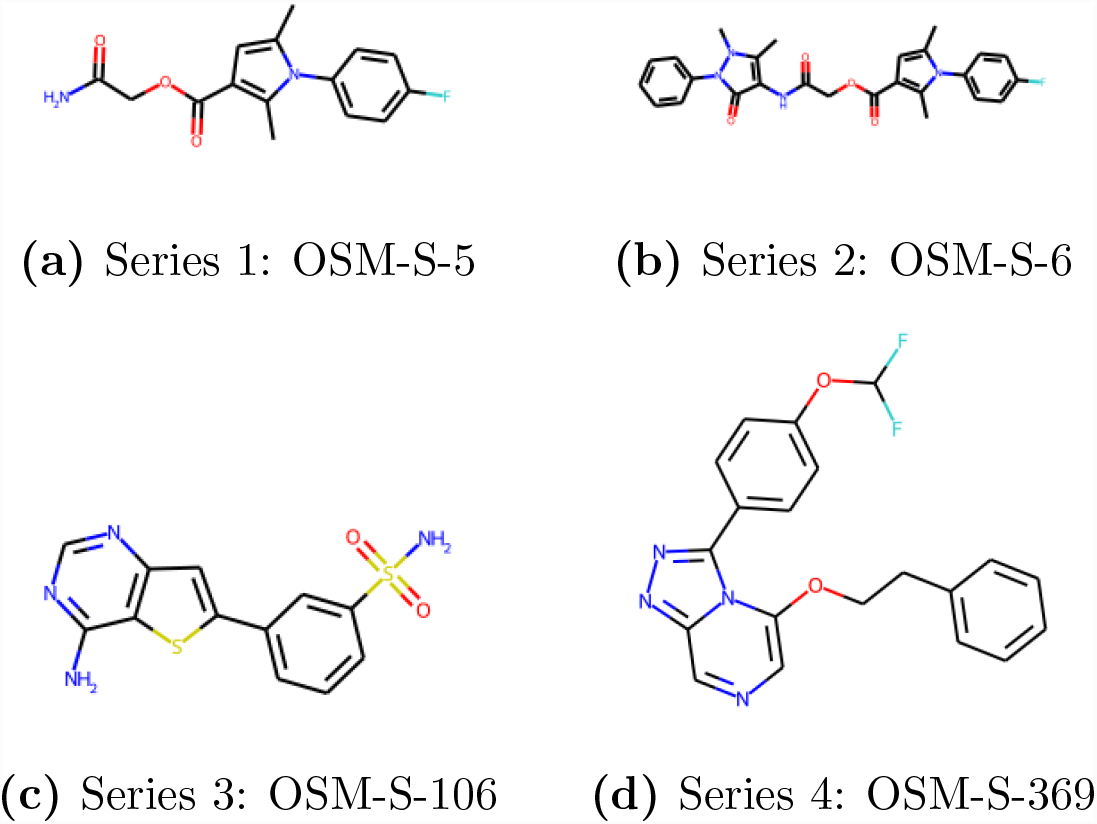
Initial compounds of each OSM series

Although targets of Series 1-3 compounds remain unknown, a promising biological target of *Plasmodium falciparum* has been identified in P-type ATPase PfATP4, a parasite cell membrane enzyme which exports Na+ ions and imports H+ ions. ^22,23^ Based on a previous study, Series 4 compounds are considered to target PfATP4.^24^ The first few series were derived from the Tres Cantos list of hits against *P. falciparum* released by Glaxo Smith-Kline in 2010^25^ but, although several potent drug candidates were found, structural difficulties have hindered progress.

In **Series 1**, a labile ester created stability concerns and potency of compounds decreased whenever changes were made to the central structure. **Series 2**, on the other hand, presented low solubility, ^26^ and in **Series 3**, the mechanisms of action of the initial compound are still under active investigation, as it is believed to inhibit one or more kinases. ^27,28^ However, the analogues derived and evaluated in the series have not exhibited high potency.

The last set, **Series 4**, is the current series of interest of OSM consortium. ^29^ These triazolopyrazine analogues were initially identified in a high throughput screening performed in 2013 by Pfizer and Medicines for Malaria Venture (MMV) ^30,31^ and contain many potent compounds, some of which have proven to be potent in vivo, and display many desirable physicochemical properties.^28^ A correlation has been found between molecular potency and parasite ion-regulated assay.^32^

This study focused first on building a predictive model for Series 4 analogues,^19^ but as our method can inherently distinguish structurally heterogeneous chemical sets, all series and assays were then considered for a more comprehensive analysis. A raw dataset containing all OSM compounds from Series 1 to 4 and their respective assay data was downloaded from the Master List of chemicals provided by OSM.^33^ Pre-processing (Figure 1(A)) was performed as outlined in:^34^ (1) compounds with no SMILES or Pfal values were removed; (2) molecular structures were normalised using RDkit,^35^ with salts stripped, canonical tautomer calculated, and charges normalised; and (3) data were deduplicated by recalculating each compound’s InChiKey. The final dataset included 386 unique compounds, each with a respective SMILES code and an associated binding activity (pIC50) to *Plasmodium falciparum*.

For each compound, circular molecular fingerprints were generated by RDKit using the Morgan algorithm. ^36^ In preliminary tests, we observed very similar performance of the modSAR algorithm for fingerprints produced with radius=2 (ECFP4) and radius=4 (ECFP8) parameters, so we selected the version with a larger radius. Therefore, our final configuration consists of Morgan circular fingerprints of radius=4 collapsed to 1024 bits, closely resembling the ECFP8 fingerprint algorithm commonly used in cheminformatics.^37^

### modSAR algorithm

ModSAR (Figure 1(B)-(E)) combines modularity clustering^38^ and regularised piecewise linear regression^21^ to learn the quantitative structural-activity relationships (QSAR) relationship of compound activity. ^20^ The algorithm involves two main stages: first modules of molecules that share similar structures are identified, and then each such module is modelled to derive piecewise linear equations, as demonstrated in Figure 1. These steps are described in more detail below.

Similarity among compounds is described by the pairwise Tanimoto coefficient *Tc*^39^ applied to the circular fingerprints.^37^ Pairs of compounds are connected by an edge in the network, if chemical similarity is above a threshold *Tc* ≥ *t*_*α*_, which is identified automatically by modSAR and corresponds to the value that optimises the average clustering coefficient of the network.^20,40^ Given the relavant similarity network, clustering partitions compounds in distinct modules by maximising the modularity metric.^41–44^ We are then able to explore the chemical space of each module separately, as compounds in the same module will typically share a common structural core or scaffold.

Having described the chemical similarities in the dataset in the first stage, the second stage derives the structure-activity mapping. Each of the modules found in the previous steps are modelled by independent piecewise linear regression equations using the OPLRAreg algorithm.^21,45^ For reference, details of OPLRAreg modelling is included in supporting information (Mathematical Details of OPLRAreg). One of the features is optimally selected to act as a breakpoint, effectively separating the data into n disjoint sub-groups called “regions”, each of which is then fitted by independent linear equations. OPLRAreg identifies all of these properties (i.e. optimal feature, number of regions and regression coefficients) simultaneously by maximising the mean absolute error (MAE) of *pIC*_50_ value through a mathematical programming optimisation model.

The algorithm derives an optimal subset of features to be used in each equation, as controlled by a regularisation parameter *λ* ≥ 0. For *λ* = 0, no regularisation is enforced and the linear equation can have as many features as possible but incurs a risk of overfitting the data. Larger *λ* values reduce the number of features included in the equation while reducing the risk of overfitting. The most common scaffold in relevant groups of compounds was identified by the rdScaffoldNetwork algorithm.^46^

### Model Inference

Piecewise linear regression equations identified by modSAR can lead to structure-activity interpretation. It is note that, as regression equations are fitted independently for each module, data can be further split into as many sub-clusters, i.e. regions, as required to minimise regression error.^20,21^ In practice, as the algorithm selects a single feature to serve as breakpoint for defining regions and as we are handling binary data, there can be at most two disjoint regions for each module.

An advantage of our methodology and the associated use of circular fingerprints, is that one can reverse each fingerprint bit to the relevant chemical fragment. As certain bits are selected by the optimisation procedure implemented in modSAR and incorporated to the regression equation, a latent association of the relevant substructure to the binding activity can be put forward. We take advantage of the bit-fragment relationship to evaluate the presence and prevalence of certain fragments in network modules and piecewise regions so as to hypothesise on their contribution to the activity of compounds.

Additionally, SHapley Additive exPlanations (SHAP) ^47^ value analysis is used to further inspect the importance of these fragment contributions.^48^ SHAP values interpret the output of a machine learning model by connecting optimal credit allocation with local explanations using the classic Shapley values from game theory. In this work, we applied and extended the work from the SHAP barplot^49^ for computing feature importance.

## Model Tuning

To identify a suitable hyper parameter setting, five-fold cross validation was performed for different *λ* values. For each *λ*, the dataset was split into five sets, each set was then used separately as test set while the other four portions were used as training set in each round of model training. The mean Root Mean Squared Error(RMSE) of the five-fold cross validation for each *λ* was utilised as evaluation metric to provide an indication of model fitness.^50^

## Results

### Cross Validation

The result of five-fold cross validation at 20 different *λ* is as shown in Figure S1a. The result shows that the RMSE in test set is almost always slightly higher than train set, at around 0.9, indicating that modSAR fits the OSM dataset well with no indication of overfitting. Overall, modSAR model performs best around *λ* = 0.06. Similar results were found using the Mean Absolute Error (MAE) metric, shown in Figure S1b and average running time for each parameter is shown in Figure S1c.

### Network modules

The chemical similarity network of the drug dataset as partitioned into clusters, is shown in Figure 3. Edges represent pairwise Tanimoto similarity that exceed the optimal threshold, which was calculated by the algorithm to be *t*_*α*_ ≥ 0.20. Nodes are coloured according to their cluster membership.

**Figure 3:**
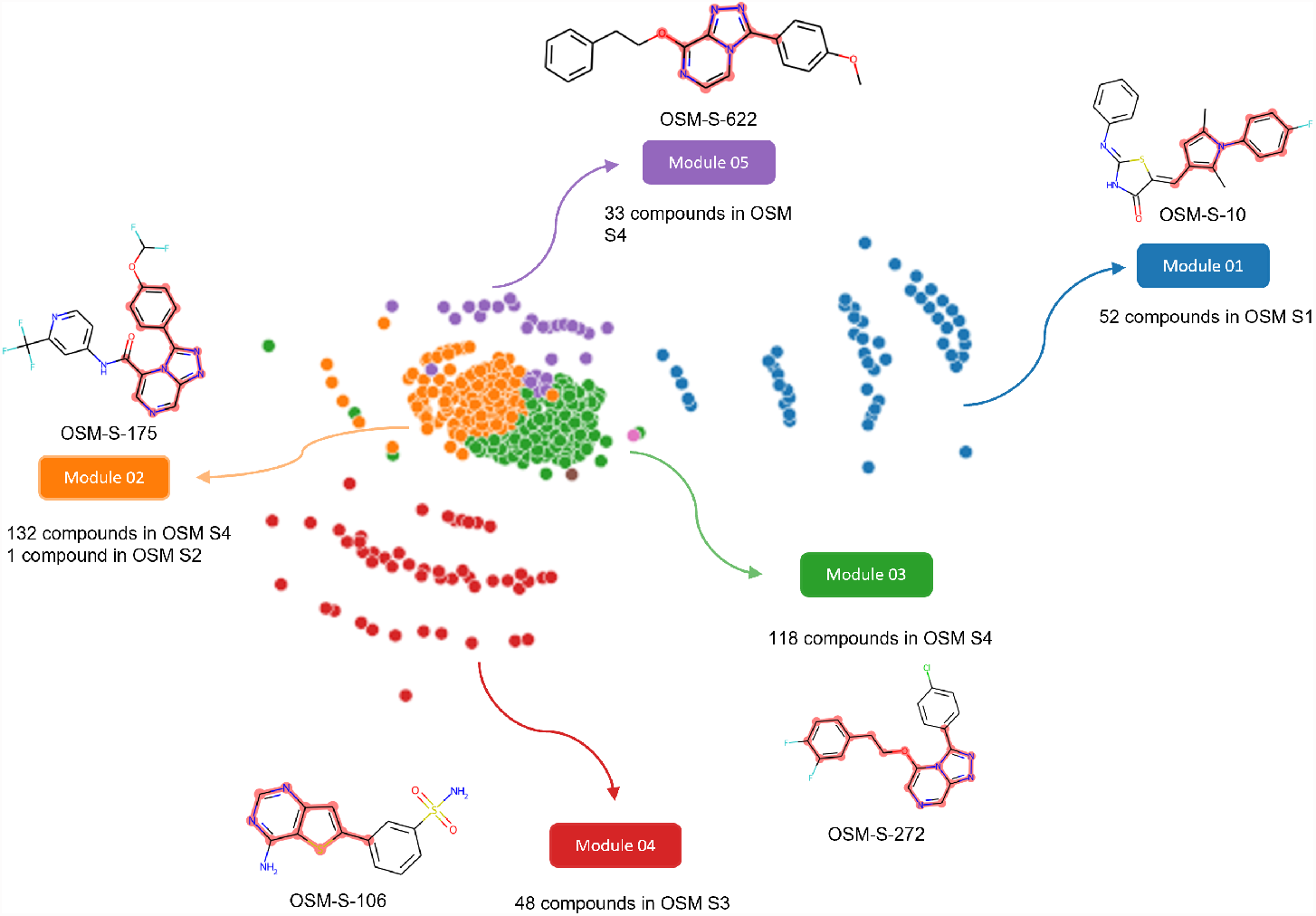
Modules identified by modularity optimisation implemented in modSAR with edges indicating Tanimoto similarity, *t*_*α*_ ≥0.20. Colours signify cluster membership. Representative compounds and dominant scaffolds for each cluster are also shown.

To provide a first visual inspection of structure-activity relationships present, the node (compound) with the highest within-module degree is selected as the representative compound of each module, thus depicting the most general structural characteristic of the neighbouring compounds. The most common scaffold in each representative compound for each module is highlighted in red (Figure 3). Note that the highlighted scaffold is the most dominant structure and not all compounds in the module may contain that substructure.

The clustering procedure represents the chemical properties of the dataset well and reflects the heterogeneity inherent in the various compound series. The five modules as partitioned by algorithm, closely match the analogue series present in OSM dataset, as shown in Table S1 and in the comparative visualisation in Figure S2. OSM Series 1 and Series 3 compounds are members of Modules m01 and m04, respectively, while Series 4 compounds have been allocated into three distinct modules: m02, m03 and m05. The Series 2 structure (OSM-S-66) was assigned to m02, as it shares structural similarities to two compounds in that module, namely OSM-S-359 and OSM-S-570. There were also two singletons, OSM-S- 89 and OSM-S-69 (not represented in the figure) as these two structures differ from the rest of the dataset.

A closer inspection of modules related to OSM Series 4 (m02, m03 and m05) and their associated highlighted scaffold in Figure 3 allows for further insight into this dataset. Each module represents core substructures that are more specific than the Series 4 triazolopyrazine core, and the assumption here is that each detected module would have their structureactivity relationships modelled individually. Therefore, the subsequent sections describe how the predicted equations of each of these modules compare, and point to molecular fragments that relate to bioactivity within the core represented by each module.

### Analysis of Modules via Regression Equations

A summary of rules and equations is shown in Table 1, where the breakpoint features and the equations identified for each cluster are shown. The distribution of *pIC*_50_ under different subsets and the presence of certain bits can be seen in Figure S3. All bits selected by modSAR are visualized in Figures S4. A detailed description and interpretation of these results follow.

**Table 1:**
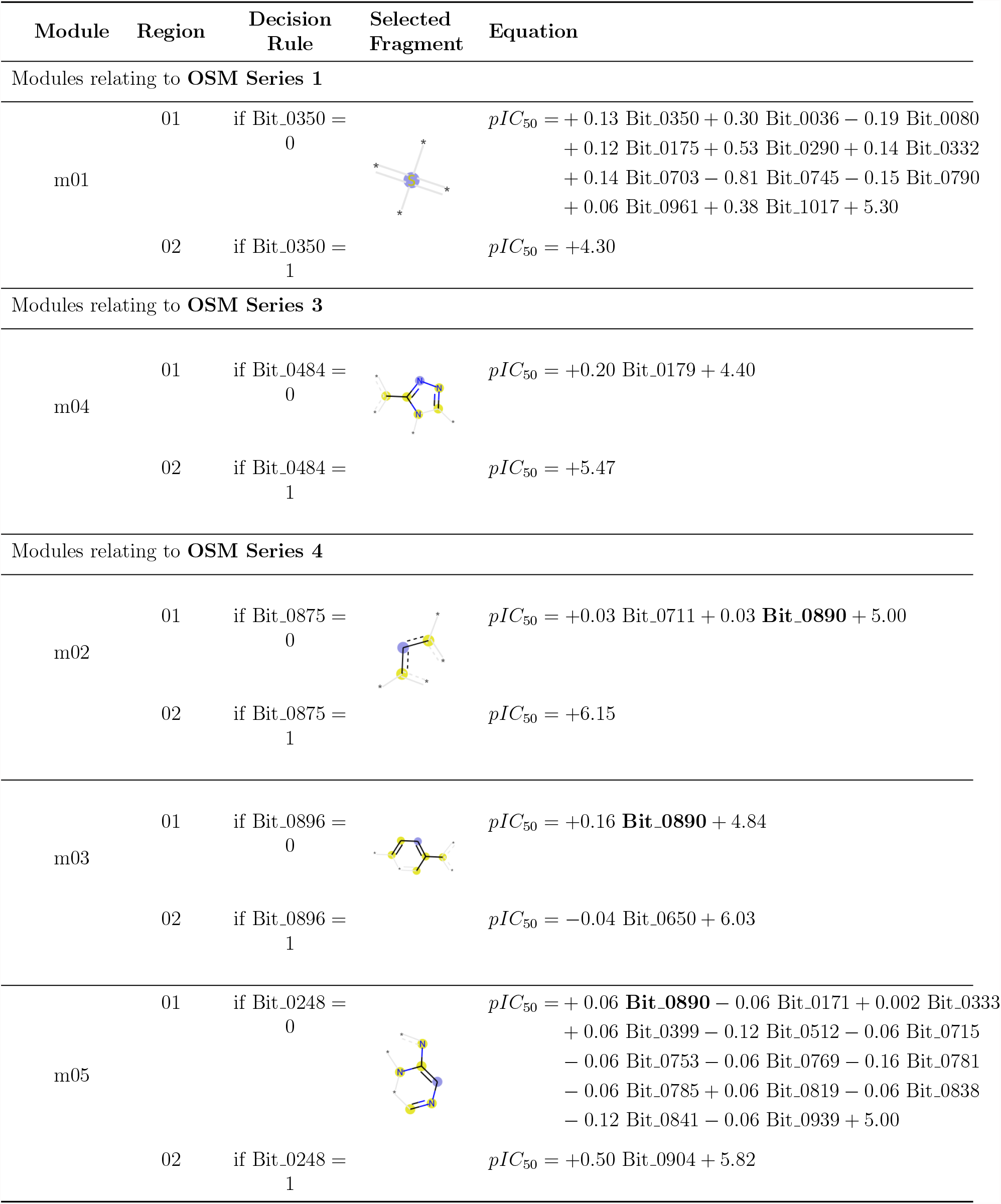
Equations and breakpoints identified for modules indicated in Figure 3.

| Module | Region | Decision Rule | Selected Fragment | Equation |
| --- | --- | --- | --- | --- |
| Modules relating to **OSM Series 1** |  |  |  |  |
| m01 | 01 | if Bit_0350 = 0 | | $pIC_{50} = +0.13 \text{ Bit\_0350} + 0.30 \text{ Bit\_0036} - 0.19 \text{ Bit\_0080}$<br>$+ 0.12 \text{ Bit\_0175} + 0.53 \text{ Bit\_0290} + 0.14 \text{ Bit\_0332}$<br>$+ 0.14 \text{ Bit\_0703} - 0.81 \text{ Bit\_0745} - 0.15 \text{ Bit\_0790}$<br>$+ 0.06 \text{ Bit\_0961} + 0.38 \text{ Bit\_1017} + 5.30$ |
| | 02 | if Bit_0350 = 1 | | $pIC_{50} = +4.30$ |
| Modules relating to **OSM Series 3** |  |  |  |  |
| m04 | 01 | if Bit_0484 = 0 | | $pIC_{50} = +0.20 \text{ Bit\_0179} + 4.40$ |
| | 02 | if Bit_0484 = 1 | | $pIC_{50} = +5.47$ |
| Modules relating to **OSM Series 4** |  |  |  |  |
| m02 | 01 | if Bit_0875 = 0 | | $pIC_{50} = +0.03 \text{ Bit\_0711} + 0.03 \text{ Bit\_0890} + 5.00$ |
| | 02 | if Bit_0875 = 1 | | $pIC_{50} = +6.15$ |
| m03 | 01 | if Bit_0896 = 0 | | $pIC_{50} = +0.16 \text{ Bit\_0890} + 4.84$ |
| | 02 | if Bit_0896 = 1 | | $pIC_{50} = -0.04 \text{ Bit\_0650} + 6.03$ |
| m05 | 01 | if Bit_0248 = 0 | | $pIC_{50} = +0.06 \text{ Bit\_0890} - 0.06 \text{ Bit\_0171} + 0.002 \text{ Bit\_0333}$<br>$+ 0.06 \text{ Bit\_0399} - 0.12 \text{ Bit\_0512} - 0.06 \text{ Bit\_0715}$<br>$- 0.06 \text{ Bit\_0753} - 0.06 \text{ Bit\_0769} - 0.16 \text{ Bit\_0781}$<br>$- 0.06 \text{ Bit\_0785} + 0.06 \text{ Bit\_0819} - 0.06 \text{ Bit\_0838}$<br>$- 0.12 \text{ Bit\_0841} - 0.06 \text{ Bit\_0939} + 5.00$ |
| | 02 | if Bit_0248 = 1 | | $pIC_{50} = +0.50 \text{ Bit\_0904} + 5.82$ |

#### Module related to OSM Series 1

The structure-activity relationship of compounds in OSM Series 1 as found by modSAR is represented by Module m01 and the full equations for this module are shown in Table 1. Most chemical compounds in this module have a common scaffold, as would be expected, with 43 of the 52 compounds containing the fragment highlighted in Figure 3.

In the regression equations, activity of compounds in this module is predicted by one of two linear equations according to the presence or absence of fragment Bit 0350. When the fragment corresponding to Bit 0350 is not present in the compound (i.e. Bit 0350 = 0), the activity of that compound is predicted by the presence of eleven fragments (Bit 0350 included). On the other hand, if a compound includes this fragment (Bit 0350 = 1), the model predicts that its bioactivity will be *pIC*_50_ = 4.30, thus inactive against *Plasmodium Falciparum*, if an activity threshold *pIC*_50_ ≥ 5.80 is assumed.^19^ By exploring **m01** (Region 01) equation, we can also see which of the remaining fragments selected by the algorithm make positive or negative contributions to the bioactivity of these compounds.

Beyond the observation of signal and magnitude of regression coefficients, we have ranked the importance of fingerprint bits according to their relative SHAP values (Figure S5a). In decreasing order of importance, the presence of fragments Bit 0290, Bit 0036, Bit 0703, Bit 0332, Bit 0031, Bit 0175, and Bit 0961 are predicted to make a positive contribution, while Bit 0745, Bit 0350, Bit 0080, Bit 1017, and Bit 0790 have a negative contribution to activity. Indeed, we can observe the association of the most positive or negative bits, Bit 0290 and Bit 0745 respectively, in the *pIC*_50_ activity of compounds (see Figure 4 and Figure S3a). Activity of compounds in this module which only contain Bit 0290 is much higher compared to those which only contain Bit 0745. The combination of positive and negative contributing fragments can be seen in Figures S6a and S6b.

**Figure 4:**
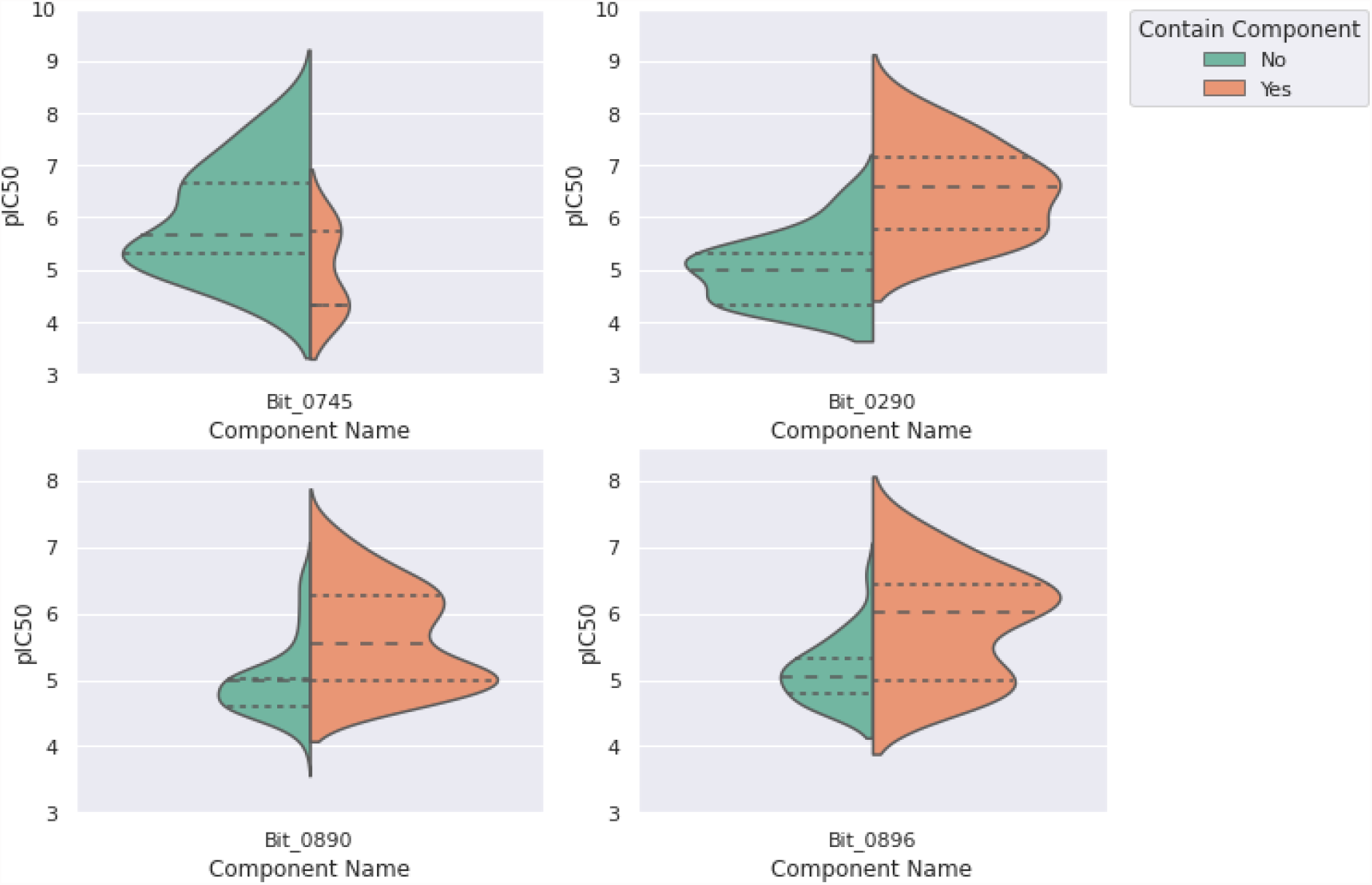
Comparison of *pIC*_50_ distributions in different subsets of the data split according to fragment presence.

### Modules related to OSM Series 3

Module **m04** describes the binding activity of compounds in OSM Series 3. The piecewise equations in Table 1 suggest that the Series 3 compounds are not likely to be active, as the maximum activity values calculated from the two equations are either *pIC*_50_ = 5.47 (when Bit 0484 = 1) or *pIC*_50_ = 4.60 (when Bit 0484 = 0 and Bit 0179 = 1), which is lower than our defined threshold of *pIC*_50_ ≥ 5.80.^19^ This is also supported by the distribution of the true *pIC*_50_ values of Series 3 compounds. As shown in Figure S3c, the median and mode value of Series 3 are both less than 5, and only 2 out of 49 compounds are considered to be active, namely OSM-S-106 and OSM-S-590. Ranking importance of the two bits can be seen in Figure S5d and combination of positive and negative contributing fragments can be seen in Figure S6g and S6h.

### Modules related to OSM Series 4

Module **m02**, Module **m03** and Module **m05** describe the binding activity of Series 2 and Series 4, as the piecewise equations demonstrated in Table 1. Since Series 2 correspond to a single compound, we focus our analysis on Series 4 properties.

One common feature of these modules is the prominence of Bit 0890 in the equations. The presence of the fragment represented by this bit is predicted to make a positive contribution towards the binding affinity of compounds. Interestingly, the structure of Bit 0890 (Figure 5b) is a close, albeit not exact, match to the triazolopyrazine core of the series (Figure 5a). If we combine all fragment bits that are predicted to make positive contributions to the binding activity of these compounds, we arrive at the fragment shown in Figure 5d. This visualisation suggests that on top of the triazolopyrazine core for Series 4, which could be approximately represented by Bit 0890 = 1 in the regression equations, the northwestern fragment should be retained Bit 0896 = 1 (Figure 5c) in order to maximise the activity of Series 4 compounds as suggested by the regression equations and the SHAP value analysis.

**Figure 5:**
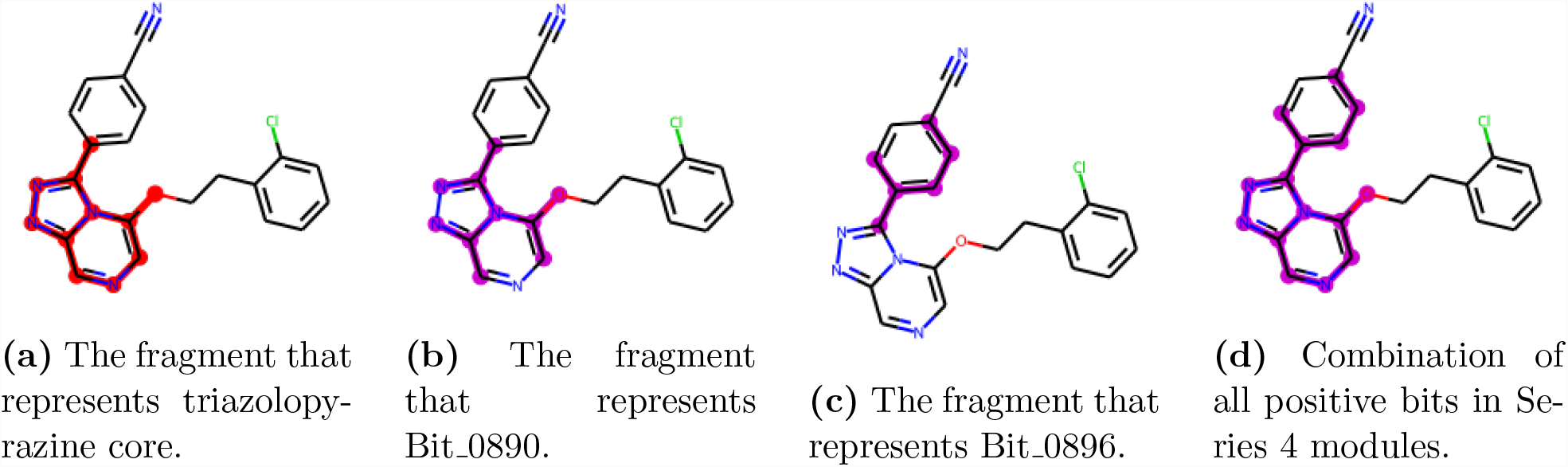
Visualisation of relevant fragments present in OSM Series 4 modules.

A similar conclusion can be drawn by comparing the two distributions *pIC*_50_ of compounds with and without Bit 0896 using a one-sided t-test with the following hypothesis:

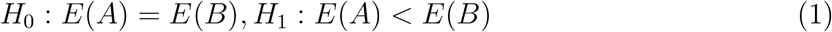

where A denotes the population of compounds which do not contain Bit 0896, B denotes the population of compounds which contain Bit 0896. The obtained *p* − *value* = 2.83*e* − 16 suggested that the null hypothesis was rejected with a confidence level at 99.9%.

A similar analysis can be made for each module. For example, Figures S6i and S6jbelow compare the positive and negative contributing fragments specific to Module m05. Additional plots about OSM Series 4 modules can be seen in Figures S5b, S5c, S5e, and S6.

### Model Evaluation

Having tuned the *λ* parameters in Model Tuning, we now assess the performance of the methodology through **Y-Randomisation** and **Applicability Domain**.

#### Y-Randomisation

Y-randomisation is employed as validation tool to compare the performance of a QSAR model with the pseudo-random models trained on permuted datasets.^51^ To verify that modSAR does not make predictions by chance, we designed the pseudo-random models as follows. Three different sets of pseudo-random data were generated via randomised fingerprints (rx), randomised *pIC*_50_ (ry), and permuted *pIC*_50_ (py). The rx was generated by assigning 0 or 1 to the 1024 fingerprints bit for each molecule, the ry was generated using random number within the range of real *pIC*_50_ value while py was generated by shuffling the real *pIC*_50_ value.

Five pseudo-random models were trained with synthetic datasets, i.e. (1) model 1: trained with rx and y; (2) model 2: trained with x and py; (3) model 3: trained with x and ry; (4) model 4: trained with rx and ry; and (5) model 5: trained with rx and py. We compared the performance of the five pseudo-random models with the original modSAR model using different *λ* (from 0.05 to 0.1) on a 10-fold cross validation. The mean and standard deviation of the model RMSE outperforms the pseudo-random datasets (Table 2). Figure S7 provides an intuitive comparison between different pseudo-random model and the original model.

**Table 2:** Comparison of test set error in y-randomization. Results shown in terms of average RMSE metric and standard deviation of the original model and permutations.

| $\lambda$ | model 1<br>y vs rx | model 2<br>py vs x | model 3<br>ry vs x | model 4<br>ry vs rx | model 5<br>py vs rx | original<br>model |
| --- | --- | --- | --- | --- | --- | --- |
| 0.05 | 0.9569<br>($\pm 0.1007$) | 0.9480<br>($\pm 0.1636$) | 1.2334<br>($\pm 0.1146$) | 1.330<br>($\pm 0.1593$) | 0.9005<br>($\pm 0.0717$) | **0.8367**<br>($\pm 0.1504$) |
| 0.06 | 0.9515<br>($\pm 0.0959$) | 0.9685<br>($\pm 0.1429$) | 1.1473<br>($\pm 0.1331$) | 1.2064<br>($\pm 0.1340$) | 1.0363<br>($\pm 0.1463$) | **0.8065**<br>($\pm 0.1302$) |
| 0.07 | 0.9092<br>($\pm 0.0732$) | 0.9093<br>($\pm 0.0811$) | 1.1387<br>($\pm 0.14284$) | 1.2959<br>($\pm 0.1399$) | 0.9208<br>($\pm 0.0639$) | **0.8660**<br>($\pm 0.1741$) |
| 0.08 | 0.9044<br>($\pm 0.1279$) | 0.9192<br>($\pm 0.1035$) | 1.1820<br>($\pm 0.0851$) | 1.2859<br>($\pm 0.0953$) | 0.8950<br>($\pm 0.1135$) | **0.8591**<br>($\pm 0.1442$) |
| 0.09 | 0.8858<br>($\pm 0.1011$) | 0.9036<br>($\pm 0.1208$) | 1.1464<br>($\pm 0.0552$) | 1.2668<br>($\pm 0.7561$) | 0.9164<br>($\pm 0.1289$) | **0.8790**<br>($\pm 0.1193$) |
| 0.10 | 0.9056<br>($\pm 0.1073$) | 0.9308<br>($\pm 0.0781$) | 1.2048<br>($\pm 0.1262$) | 1.2224<br>($\pm 0.0936$) | 0.8908<br>($\pm 0.0864$) | **0.8819**<br>($\pm 0.1462$) |

#### Applicability Domain

The applicability domain (AD) defines the chemical space covered by the model, indicating the reliability of the model in predicting new compound properties. In this study, the AD of modSAR is determined by the leverage approach^50^ which calculates the leverage and standard residual of a compound, and visualize all the compounds in a Williams plot. A critical leverage value h* is calculated by the equation: *h*^*^ = 3*p*′*/n*, where *p*′ is the number of model variables plus one, and n is the number of the objects used to calculate the model. To reduce the computation complexity caused by the high dimension of data, the dimension was reduced to two using Principal Components Analysis (PCA) before calculating the AD. As shown in Figure 6, all compounds were inside the applicability domain.

**Figure 6:**
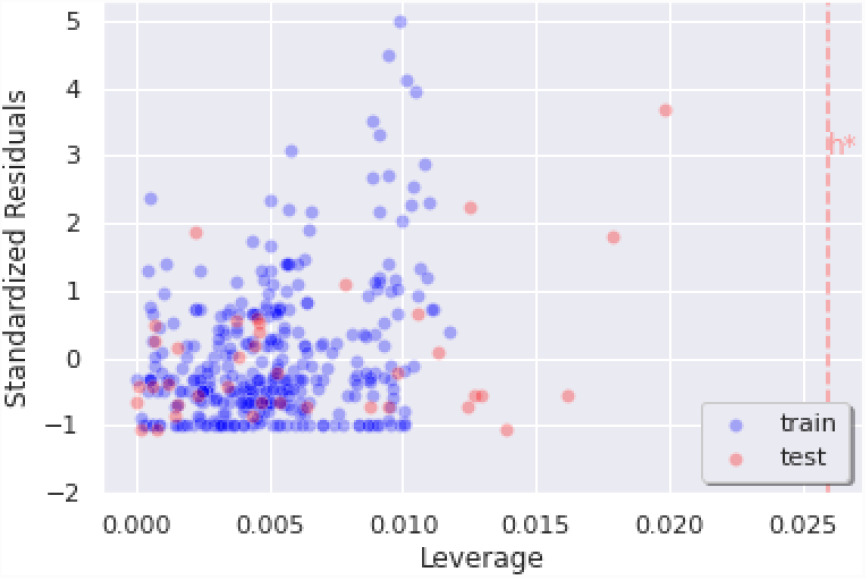
Williams Plot to evaluate the applicability domain of modSAR

#### Virtual Screening

To identify potential molecules with antimalarial properties, virtual screening was performed on ChEMBL^52^ database entries. As shown in Figure S8, all compounds within the defined applicability domain which contain any max common pattern of each module were selected, and then their activity was predicted using the piecewise equations for each module. In total, 7569 compounds from ChEMBL database fall in the neighborhood of OSM dataset defined by modSAR. We excluded 48 compounds that share the same canonical SMILES with the compounds from the OSM dataset, and computed the activity of the remaining 7521 compounds. Given the threshold of pIC50 greater than 5.8, 279 compounds are predicted to be active against malaria, which includes 55 compounds which are already selected as candidate compounds for malaria by ChEMBL through open source malaria screening. Finally, the remaining 224 compounds are novel predictions by modSAR, as listed in supporting file ‘Prioritized Molecules.xlsx’.

Given the 224 novel predictions, we looked into fragments prioritization for Series 4 as it is the most promising series suggested by OSM. We first re-allocated the novel compounds to each module according to their modSAR prediction. Next, we selected the compounds containing the triazolopyrazine core as well as the fragment Bit 0896 suggested by modSAR inference. In the end, three most promising compounds from the prioritised candidates were selected, as shown in Figure 7.

**Figure 7:**
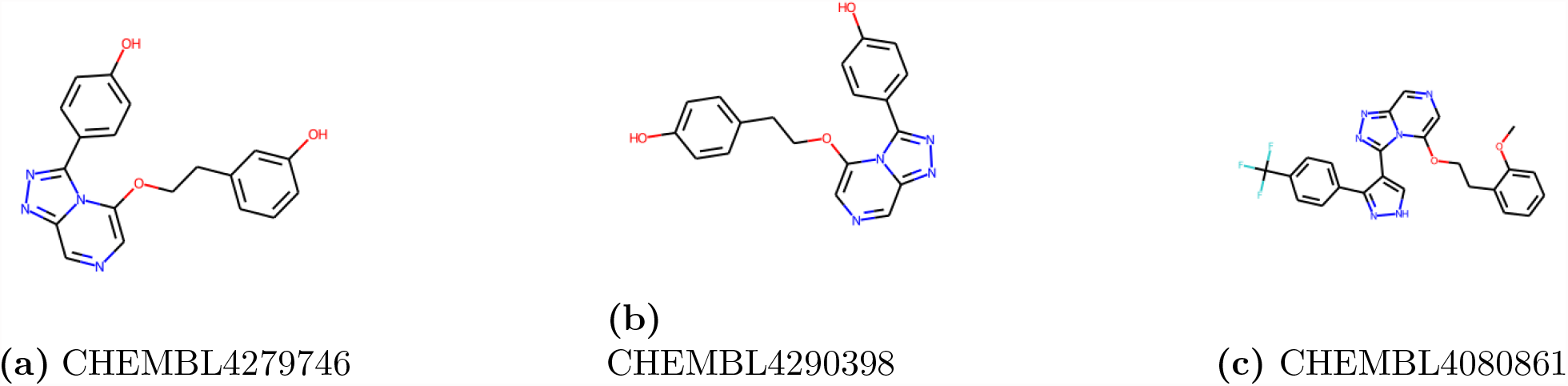
Potential antimalarial drug candidates identified via virtual screening

## Conclusion

Computational methodologies for target prediction^10^ and drug discovery have significant potential in malaria research. Here, we report the use of optimisation-based regression modelling coupled with network clustering in mining and analysing data related to antimalarial molecules. We illustrated use of the modSAR piecewise linear regression model method applied in the OSM dataset with ECFP fingerprints as features to describe each compound. Analysis of results showed that the method reflected the heterogeneity among different series in OSM and that it was capable of modulating separate piecewise linear equations for each molecule group. Finally performance was assessed by cross validation, randomisation and applicability domain tests, with results indicating promising performance and interpretable outputs.

An important aspect of this work lies in the mathematical nature of the modSAR model that offers explainable output, as molecular fingerprint bits selected by each equation can be reversed back to identical molecular fragments, and thus provide insights towards drug discovery. In this work we have paid particular attention in demonstrating the ability of modSAR to provide insights of datasets and prioritise useful chemical fragments. A barrier in accurate activity prediction may be that the method only selects a subset of binary fingerprints, thus only predicting discrete values, so future work will focus on combining continuous descriptors with fingerprints to improve the prediction performance.

## Supporting information

Prioritized Molecules

## Data and Software Availability

We provide all files for reviewers’ reference, and we will make our code publicly available after acceptance.

## Supporting Information Available

The Supporting Information is available free of charge in supporting information.pdf and Prioritized Molecules.xlsx.

Mathematical details of OPLRAreg, equivalency between modSAR modules and OSM series, average model performance of cross validation, pIC50 distribution of compounds related to the important bits, the visualisation and SHAP value of each important bit, model performance compared with pseudo random models, and virtual screening procedures (supporting information.pdf).

ChEMBL ID and SMILES of prioritised compounds (Prioritized Molecules.xlsx).

## Acknowledgement

We acknowledge the open and collaborative research effort of Open Source Malaria without which this study would not have been possible. Yutong Li is supported by the China Scholarship Council. Lazaros G. Papageorgiou acknowledges funding from EPSRC (EP/V01479X/1)

