## Supporting information for "Optimisation-based modelling for drug discovery in malaria"

### Mathematical Details of OPLRAreg

The mathematical model for piecewise linear regression based on mathematical programming has been reported in[1] and then refined in.[2]

Below, the details of the mathematical model are provided for reference. The objective function of OPLRAreg is shown in Equation S1 below,

$$z = MAE + \lambda \cdot REG \tag{S1}$$

where $\lambda$ is a positive user-defined parameter that controls the influence of regularisation. Variables $MAE$ and $REG$ are defined by the set of equations below,

$$MAE = \frac{\sum_s E_s}{|s|} \tag{S2}$$

$$REG = \sum_f W_{rf}^+ \tag{S3}$$

$$W_{rf}^+ \geq W_{rf} \qquad \forall r, f \tag{S4}$$

$$W_{rf}^+ \geq -W_{rf} \qquad \forall r, f \tag{S5}$$

where $E_s$ indicates the absolute error for each sample $s$ and $|s|$ is the number of samples in the training set. Positive variables $W_{rf}^+$ are introduced to indicate the absolute value of regression coefficients $W_{rf}$ and are defined by the two auxiliary constraints S4 and S5.

At each iteration, the number of regions $R$ and the partition feature $f^*$ used to identify breakpoints are fixed. The allocation of samples to regions $r \in 1, 2, ..., R$ is modelled with binary variables $F_{sr}$ while the breakpoints are represented by the free variables $X_{rf}$, where $f$ always corresponds to the partition feature $f^*$ of the current iteration.

Equation S6 guarantees that a sample can belong to only one region:

$$\sum_r F_{sr} = 1 \qquad \forall s \tag{S6}$$

while Equation S7 below ensures that breakpoints are consistent:

$$X_{rf^*} \geq X_{r-1,f^*} \qquad \forall r = 2, 3, ..., R-1 \tag{S7}$$

Equations S8 and S9 assign samples to the correct regions according to the breakpoints.

$$A_{sf^*} \geq X_{r-1,f^*} - U(1 - F_{sr}) \qquad \forall s, r = 2, 3, ..., R \tag{S8}$$

$$A_{sf^*} \leq X_{rf^*} + U(1 - F_{sr}) \qquad \forall s, r = 1, 2, ..., R-1 \tag{S9}$$

The predicted value $P_{sr}$ for sample $s$ in region $r$ is computed by Equation S10, according to regression coefficients $W_{rf}$ and the intercept $B_r$ for each region. Equation S11 and S12 compute the absolute error in prediction $E_s$ for each sample. $O_s$ are the observed values for sample $s$ and $U$ is a large number that will force these constraints to consider only the predicted values $P_{sr}$, where sample $s$ belongs to region $r$, $F_{sr} = 1$.

$$P_{sr} = \left( \sum_{f} W_{rf} A_{sf} \right) + B_r \qquad \forall s, r \tag{S10}$$

$$E_s \geq O_s - P_{sr} - U(1 - F_{sr}) \qquad \forall s, r \tag{S11}$$

$$E_s \geq P_{sr} - O_s - U(1 - F_{sr}) \qquad \forall s, r \tag{S12}$$

The summary of the mixed integer linear programming model, OPLRAreg, is given by:

$$minimise \quad z$$

$$subject\ to \quad \text{Equations (S1)} - \text{(S12)}$$

**Table S1:** Equivalency between modules and OSM Analogues Series Data

| Modules | Series 1 | Series 2 | Series 3 | Series 4 |
|---|---|---|---|---|
| **m01** | 52 | 0 | 0 | 0 |
| **m02** | 0 | 1 | 0 | 132 |
| **m03** | 0 | 0 | 0 | 118 |
| **m04** | 0 | 0 | 48 | 0 |
| **m05** | 0 | 0 | 0 | 33 |

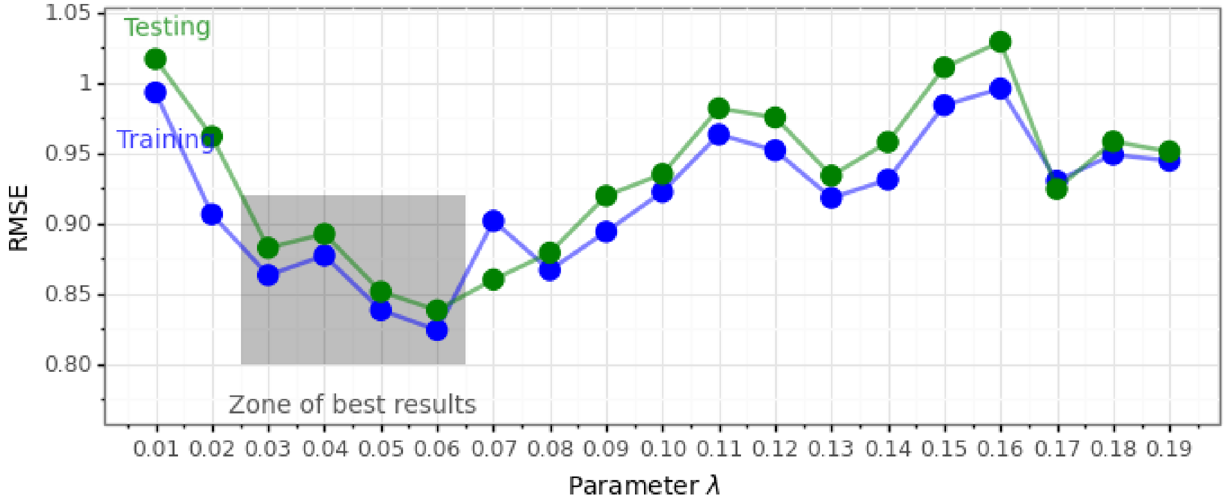


**(a)** RMSE on different $\lambda$

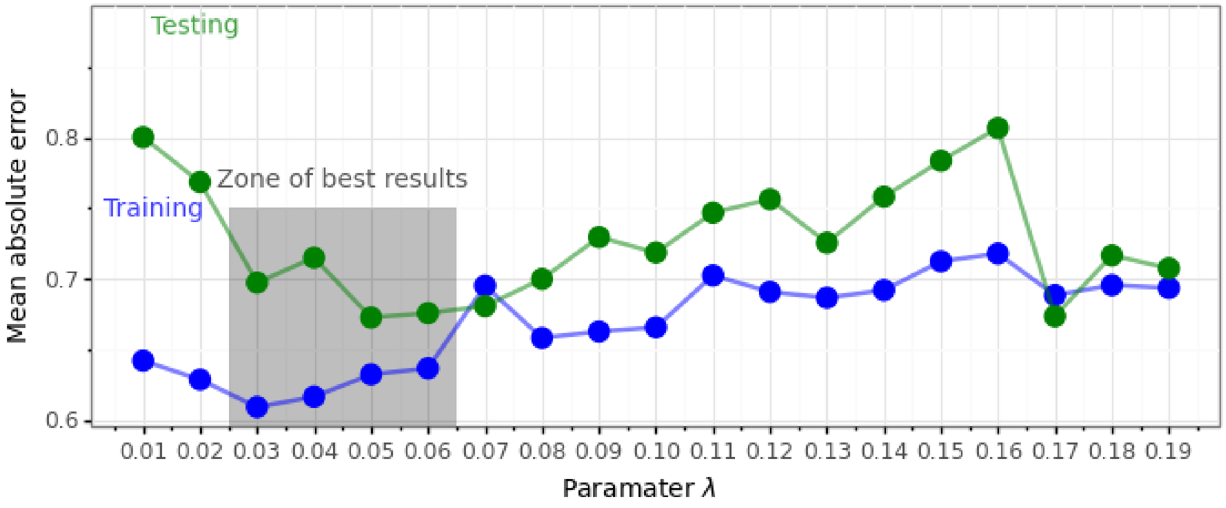


**(b)** MAE on different $\lambda$

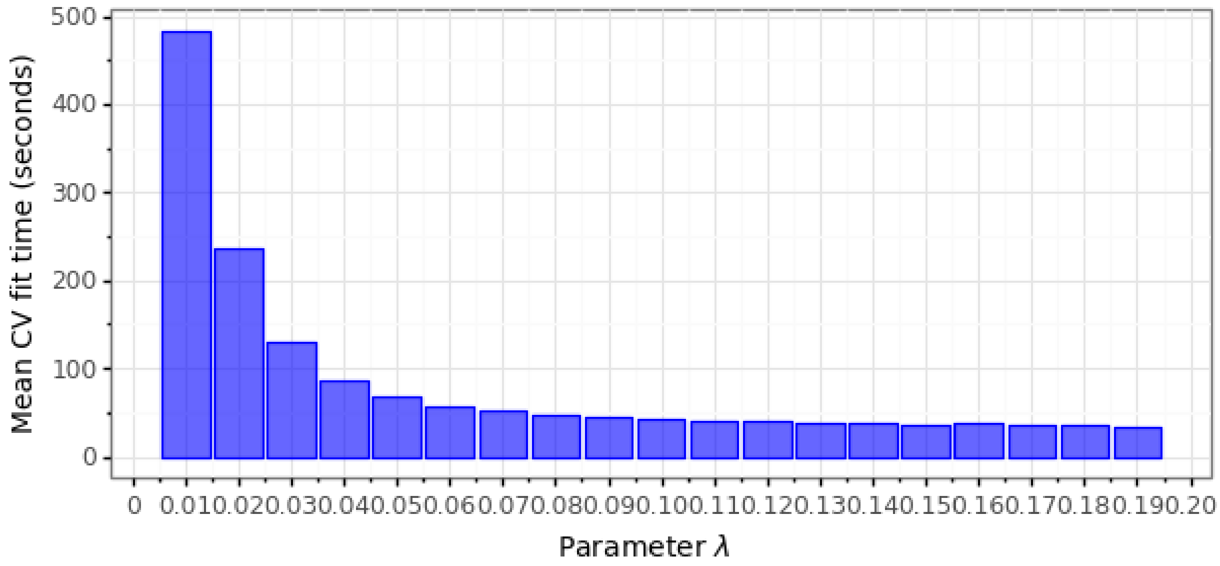


**(c)** Fit time on different $\lambda$



**Figure S1:** The average model performance of cross validation on different $\lambda$. (a) RMSE (b) MAE (c) fit time. The model achieves the best performance at $\lambda = 0.06$. And the fit

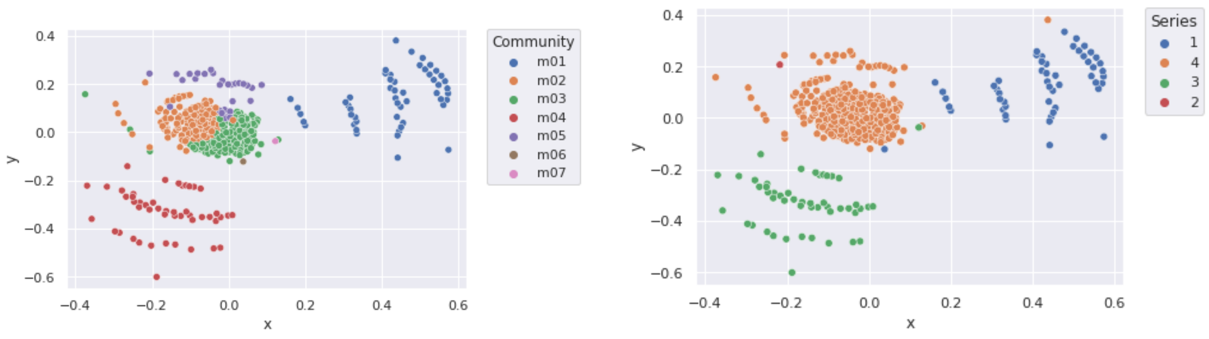


**(a)** modSAR clusters with different modules **(b)** modSAR clusters with different series

**Figure S2:** Equivalency between modules and OSM Analogues Series Data. Comparing the clusters between (a) and (b), it is easy to observed that the clusters identified by modSAR modules is corresponded to the original OSM series. ModSAR is able to detect the heterogeneity among series.

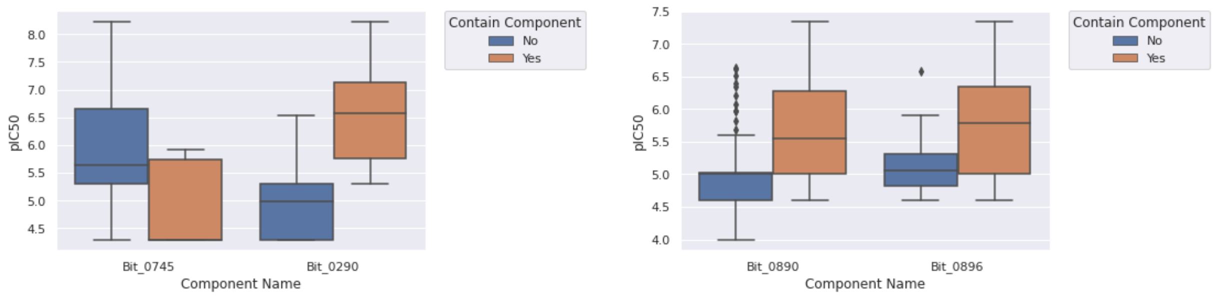


**(a)** Boxplot for Bit_0745 and Bit_0290 **(b)** Boxplot for Bit_0890 and Bit_0896

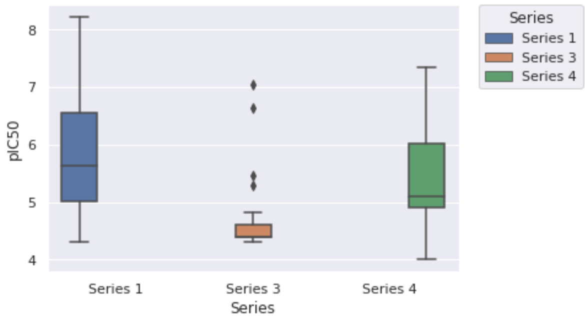


**(c)** Boxplot for each series

**Figure S3:** pIC50 value distribution of compounds related to the important bits

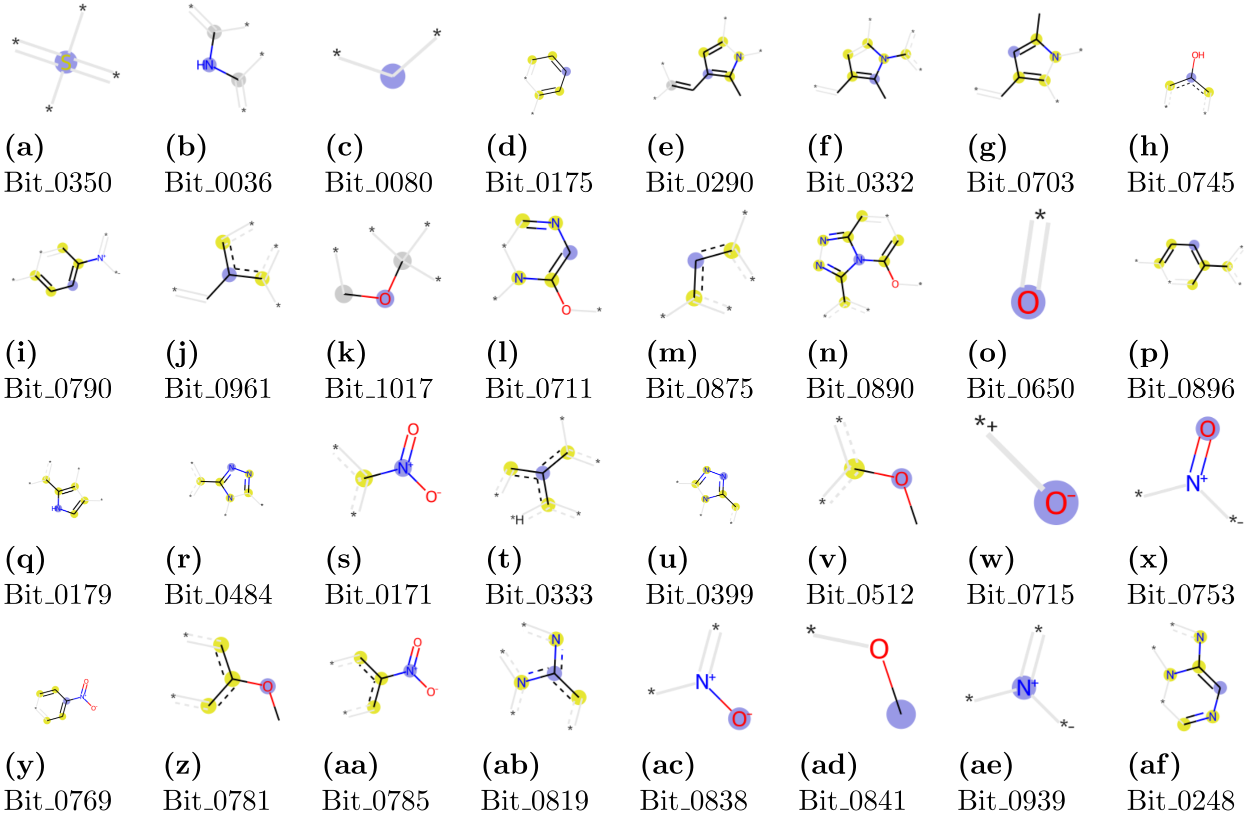


**Figure S4:** All the important fingerprint bits prioritized by modSAR and the SHAP value are shown here. (a)-(k) are the bits selected by m01; (l)-(n) are the bits selected by m02 ; (n)-(p) are the bits selected by m03; (q)-(r) are the bits selected by m04; (n), (s)-(af) are the bits selected by m05.

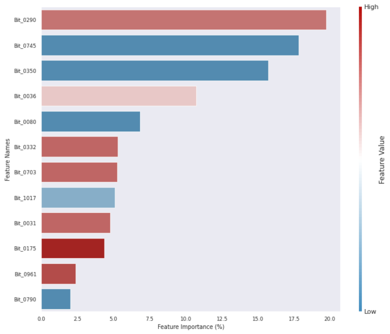

**(a)** The SHAP value of each selected bit in m01. Bit_0290, Bit_0036, Bit_0332, Bit_0703, Bit_0031, Bit_0175, and Bit_0961 make positive contributions; and Bit_0745, Bit_0350, Bit_0080, Bit_1017, and Bit_0790 make negative contributions.

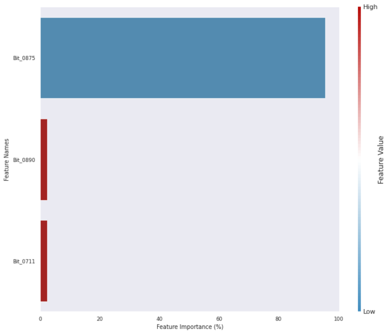

**(b)** The SHAP value of each selected bit in m02. Bit_0890, and Bit_0711 make positive contributions; and Bit_0875 make negative contributions.

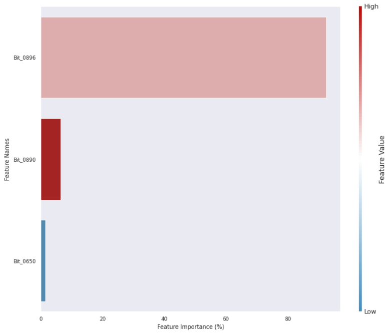

**(c)** The SHAP value of each selected bit in m03. Bit_0896, Bit_0890 make positive contributions; and Bit_0650 make negative contributions.

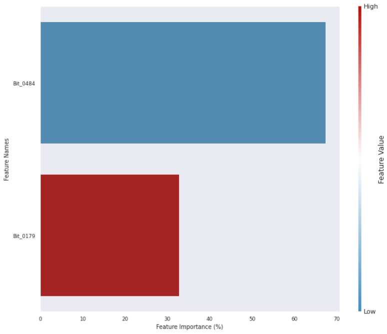

**(d)** The SHAP value of each selected bit in m04. Bit_0179 makes positive contributions; and Bit_0484 makes negative contributions.

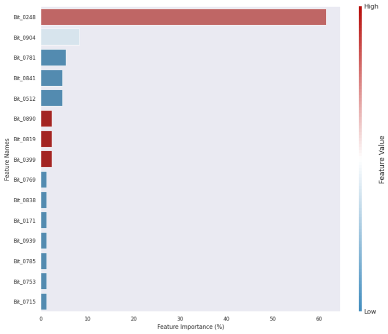

**(e)** The SHAP value of each selected bit in m05. Bit_0248, Bit_0890, Bit_0819, and Bit_0399 make positive contributions; and Bit_0904, Bit_0781, Bit_0841, Bit_0512, Bit_0769, bit_0838, Bit_0171, Bit_0939, Bit_0785, Bit_0753, and Bit_0715 make negative contributions.

**Figure S5:** The SHAP value of each selected bit.

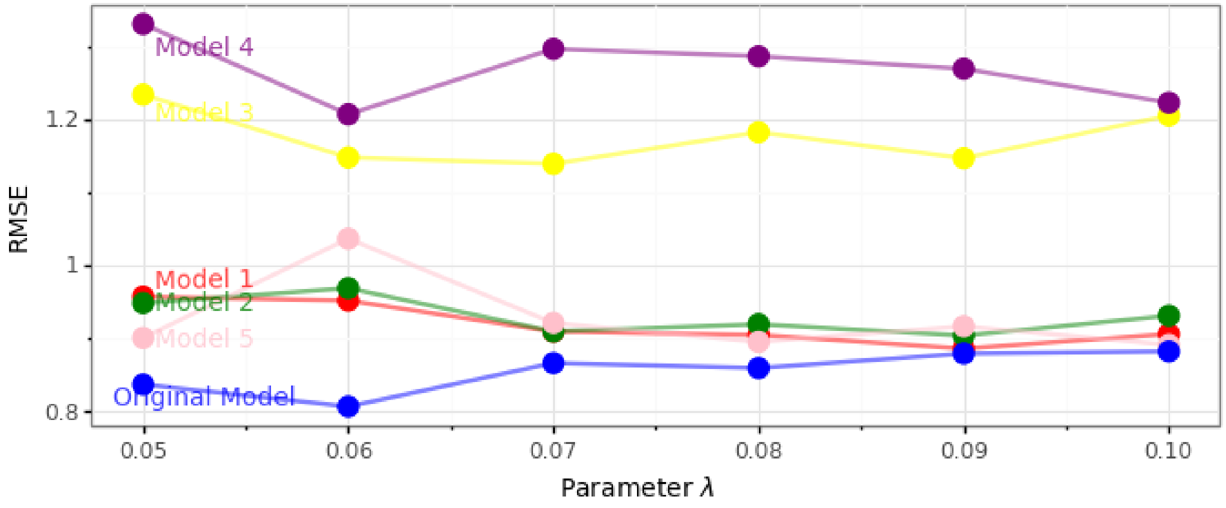


**Figure S7:** Comparisons with pseudo random models

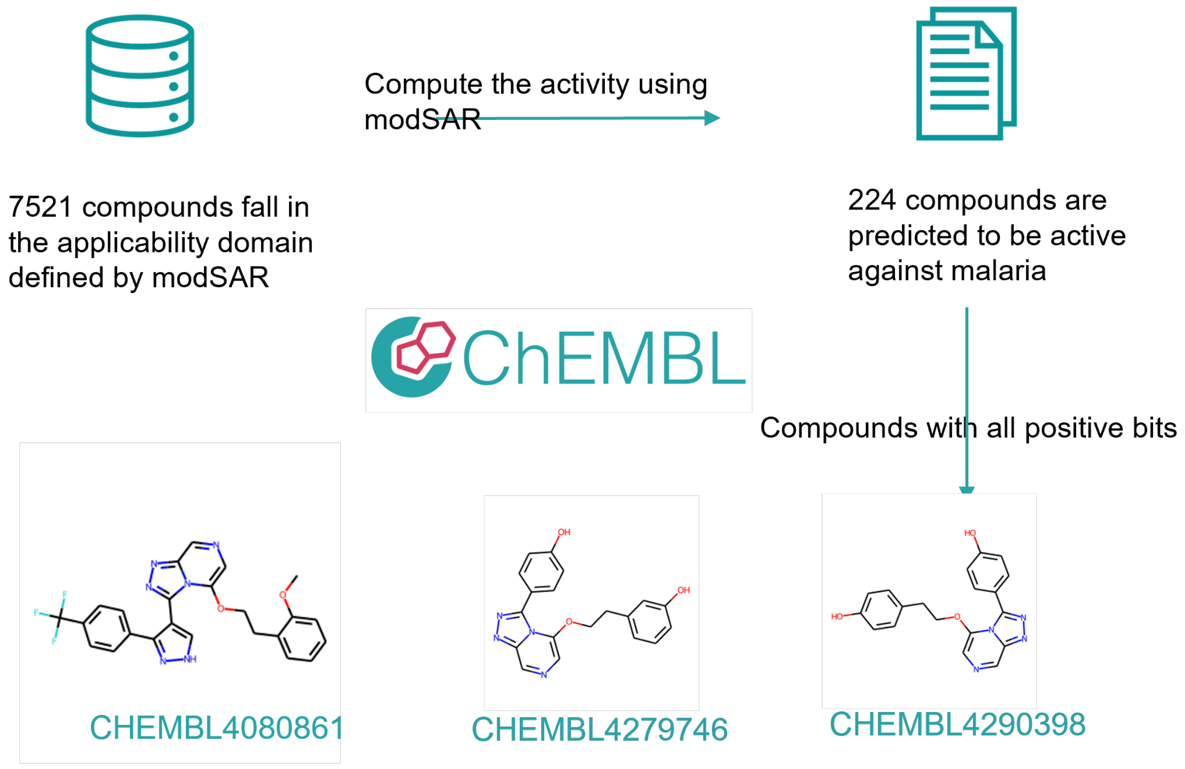


**Figure S8:** Virtual screening procedures

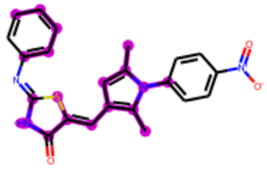

**(a)** All positive bits combined in m01

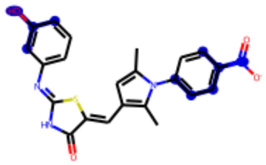

**(b)** All negative bits combined in m01. Note that because there is no compound in m01 which contains all the negative bits, here we only present Bit_0745 and Bit_0790.

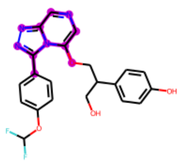

**(c)** All positive bits combined in m02

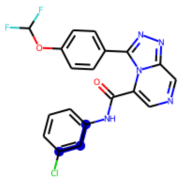

**(d)** All negative bits combined in m02

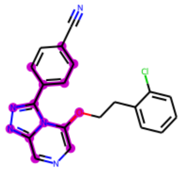

**(e)** All positive bits combined in m03

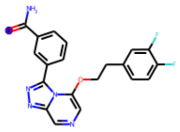

**(f)** All negative bits combined in m03

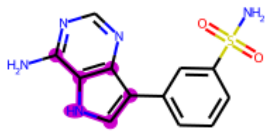

**(g)** All positive bits combined in m04

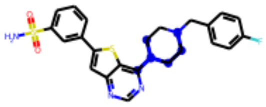

**(h)** All negative bits combined in m04

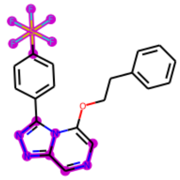

**(i)** All positive bits combined in m05

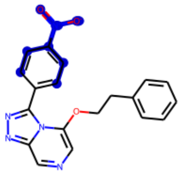

**(j)** All negative bits combined in m05

**Figure S6:** Visualisation of combined bits which make positive contribution and negative contribution in molecules
